# MdHY5 and MdHY5S form a positive transcription loop to regulate browning and phenolics synthesis in fresh-cut apple under purple LED light

**DOI:** 10.1101/2023.11.13.566910

**Authors:** Juntong Jin, Liyong Qi, Shurong Shen, Shuran Yang, Hui Yuan, Aide Wang

## Abstract

Enzymatic browning significantly affects the appearance and quality of fresh-cut fruit. Light treatment can effectively inhibit fresh-cut apple browning via unknown molecular mechanisms. Here, we found that the application of purple LED light decreased the browning index of fresh-cut apple, delaying browning as compared to that of fresh-cut apple placed in the dark, and suggesting that purple LED light suppresses browning. In addition, the expression levels of *MdHY5* and *MdHY5S*, important BASIC LEUCINE ZIPPER DOMAIN (bZIP) transcription factors that are involved in the light signaling pathway, were increased by purple LED light treatment. Silencing *MdHY5* and *MdHY5S* in apple meant that purple LED light treatment no longer inhibited fresh-cut apple browning, the expression levels of the browning-related genes *POLYPHENOL OXIDASE* (*MdPPO*) and *PEROXIDASE* (*MdPOD*) increased, and the expression of the phenolic synthesis gene *PHENYLALANINE AMMONIALYASE* (*MdPAL*) decreased. Further study revealed that *MdHY5* and *MdHY5S* bind to the *MdPPO* and *MdPOD* promoters, reducing their transcription. In contrast, *MdHY5* and *MdHY5S* bind to the *MdPAL* promoter, enhancing transcription. Further research revealed that MdHY5 and MdHY5S also bind directly to each other’s promoters to form a positive transcriptional loop that activates their transcription. Our findings revealed that purple LED light inhibits browning and increases phenolics synthesis in fresh-cut apple by activating *MdHY5* and *MdHY5S*. The results provide a theoretical basis upon which the new methods for improving the appearance and quality of fresh-cut apple can be based.

## INTRODUCTION

Freshly cut fruit and vegetable are popular with consumers because of their convenience, with fresh-cut fruit accounting for 29 % of the total fruit consumption in Europe and America (Yildiz et al. 2018), and the fresh-cut fruit industry in Asia increasing in popularity, according to the Food and Agriculture Organization of the United Nations (FAOSTAT). Fresh-cut fruit account for 11 % of the fruit consumed in Japan and South Korea (Sucheta et al. 2020) and while the market for fresh-cut products has increased owing to the convenience and health impacts (Ragaert et al. 2004). Physiological and biochemical problems such as browning, off-flavors, aging, softening, and deterioration occur during the production of fresh-cut fruit and vegetable, damaging appearance and quality of the fruit. Browning is a major problem in fresh-cut fruit and vegetable (Supapvanich et al. 2011), especially white-fleshed fruit such as apple and pear (Supapvanich et al. 2012; Zheng et al. 2019). Based on differences in the production mechanism, browning has been classified into ‘enzymatic browning,’ in which polyphenol oxidase (PPO) catalyzes phenolic compounds, and ‘non-enzymatic browning’, in which a series of complex reactions between reducing sugars (carbohydrates) and amino acids/proteins occurs (Manzocco et al. 2020). Enzymatic browning is the main cause of browning in fresh-cut fruit and vegetable (Koushesh Saba and Sogvar 2016), rendering its inhibition of great significance for the fresh-cut fruit and vegetable industry.

In typical plant cells, PPO is localized in cytoplasmic organelles such as chloroplasts, whereas phenolic substrates are mostly found in the vacuole (Toivonen and Brummell 2008). Enzymatic browning occurs in the presence of oxygen. Under mechanical damage such as cutting, PPO oxidizes phenolics to produce quinones, which interact with amino acids and proteins to form melanin, inducing browning (Taylor and Clydesdale 1987). Phenolics, oxygen, and enzymes are thus considered important factors that affect browning in fresh-cut fruit. Phenolics are beneficial to humans owing to their antioxidant activity, and although phenolics act as substrates during the browning process, it has been reported that browning can be inhibited by increasing the phenolic content and improving the antioxidant capacity (Li et al. 2022). Phenolics are synthesized via the phenylalanine metabolic pathway, in which phenylalanine ammonia-lyase (PAL) is the first rate-limiting enzyme. Increasing the PAL expression therefore increases the content of phenolic compounds, and PAL has been reported associated with browning in fruit and vegetables (Min et al. 2017).

Studies have shown that several enzymes are involved in enzymatic browning, with PPO and peroxidase (POD) the two main enzymes that are associated with fruit browning. PPO, a membrane protein that includes copper ions, is mostly found in the cytoplasm, although some has been observed on cell membranes or walls. PPO catalyzes the oxidation of polyphenols, leading to browning after cutting. In general, enzymatic browning in fruit and vegetable depends on the presence of PPO; thus, the enzyme is considered key enzyme in the browning process. Grape skin browning has been associated with increased *VvPPO* expression (Suehiro et al. 2014), and silencing *MdPPO* in apple has been found to prevent discoloration for some time after cutting (Stowe and Dhingra 2021). The use of small DNA inserts has been found to reduce the expression of genes coding for PPO, reducing the enzymatic browning effect in potatoes (Chi et al. 2014). POD is an oxidoreductase that affects fruit growth and development, particularly during maturation and aging, and has been reported to cause browning by participating in the oxidation of quinone, glutathione, and ascorbic acid (Wang et al. 2019). H_2_S treatment has been found to inhibit browning in fresh-cut lotus root slices by inhibiting PPO and POD activities (Sun et al. 2015). Plasma-processed air reduces the activities of both PPO and POD, preventing blackening in cut apple and potatoes (Bußler et al. 2017). These results demonstrate the significant roles that PPO, POD, and PAL play in enzymatic browning.

Currently, an increasing number of physical and chemical methods are being used to inhibit browning in fresh-cut fruit. However, drawbacks such as chemical residues and high costs that are associated with these treatments means that there is an urgent need to develop environmentally friendly and cost-effective techniques that can inhibit browning in fresh-cut production. Light is an environmental factor that involve in regulating both the growth and development of plants (Li et al. 2020), with previous studies reporting that light inhibits browning. For example, UV-C inhibits browning in button mushrooms by inhibiting the activity and gene expression of *AbPPO* (Lei et al. 2018). UV-C irradiation has also been found to inhibit browning and delay quality degradation on the surface of freshly cut lettuce (Han et al. 2021). Ultraviolet UV-A treatment delays the browning of fresh-cut apple and pear by reducing the PPO activity in fruit (Lante et al. 2016), while UV-B has been combined with MCP to effectively reduce the activities of both PPO and POD in fresh-cut peaches (Li et al. 2022). Although light has been reported to inhibit browning in fresh-cut apple, few reports have described the use of visible light to inhibit browning, and the mechanism by which light inhibits browning is not yet understood.

The HY5 transcription factors in plants belong to the basic leucine zipper (bZIP) family, with functional characteristics that are mainly focused on growth and development (Gangappa et al. 2016). For example, HY5 binds to the TZP promoter to mediate FR light-induced TZP expression (Li et al. 2022), and regulates the accumulation of tomato anthocyanins under blue light by binding to a promoter of the anthocyanin synthesis gene (Liu et al., 2018). CAM7 interacts with HY5 to regulate the development of Arabidopsis seedlings (Abbas et al. 2014). Studies show that HY5 affects metabolic pathways, and regulates the expression of PSY, the chlorophyll biosynthesis LHCA4, PORC, and GUN5, increasing carotenoid biosynthesis (Toledo-Ortiz et al. 2014). Another recent report showed that HY5 activates the FLS promoter to increase the number of anthocyanins and flavanols produced under solar UV (Henry-Kirk et al. 2018). Further analysis has demonstrated that *SlHY5* suppresses ripening, carotenoid biosynthesis, and anthocyanin accumulation (Wang et al. 2021). However, a role for HY5 in the inhibition of browning by light has not yet been reported.

Fresh-cut apple occupy an important position in the fresh-cut fruit market, and improving the appearance and quality of fresh-cut apple is important for the development of the fresh-cut industry. The most popular cultivar ‘Fuji’ is prone to browning after cutting. In this study, purple light was found to play a key role in inhibiting the browning and promoting phenolic synthesis of fresh-cut apple by inducing *MdHY5* and *MdHY5S* expression and silencing MdHY5 and MdHY5S in fruit by purple light treatment, thus increasing *MdPPO* and *MdPOD* and decreasing *MdPAL* expression. The experimental results reveal that MdHY5 and MdHY5S bind to the promoters of *MdPPO* and *MdPOD* and inhibit their transcription, thereby inhibiting browning. In addition, MdHY5 and MdHY5S bind to the *MdPAL* promoter, thus inhibiting the browning of fresh-cut fruits by increasing the phenolic synthesis. Moreover, MdHY5 and MdHY5S interact with their corresponding promoters, resulting in the formation of a positive transcriptional regulatory loop. These findings reveal the regulatory mechanism by which purple light inhibits browning and promotes phenolic synthesis in fresh-cut apple, thus providing a theoretical basis and practical method for improving the appearance and quality of freshly cut apple.

## Results

### The purple light-induced inhibition of browning in fresh-cut apple

To clarify the role of light in the browning of fresh-cut apple, the browning index (BI) of fresh-cut ‘Fuji’ fruit was measured using fruit maintained under different light qualities. The same light intensity of 700 lx was used and the experiment run over 4 d at 10 °C, with samples collected every 2 d. Fruit were harvested when apple were collected as part of the commercial harvest, 180 days after full bloom (DAFB) in 2020, and stored at 4 °C for later use. The results showed that different light qualities reduced the browning index of apple to varying degrees, with all inhibiting browning to some extent. However, purple light was found most effective in terms of suppression (Figure 1A, B; Figure S1A, B). To verify the results obtained, ‘Hanfu’ and ‘Lvshuai’ apple were also treated with light of different intensities, and were observed to brown rapidly; however, again purple light was observed to significantly decrease the browning index and inhibit browning in both ‘Hanfu’ (Figure S1C, D) and ‘Lvshuai’ apple (Figure S1E, F). These results demonstrate that purple light suppresses browning in fresh-cut apple.

**Figure 1.**
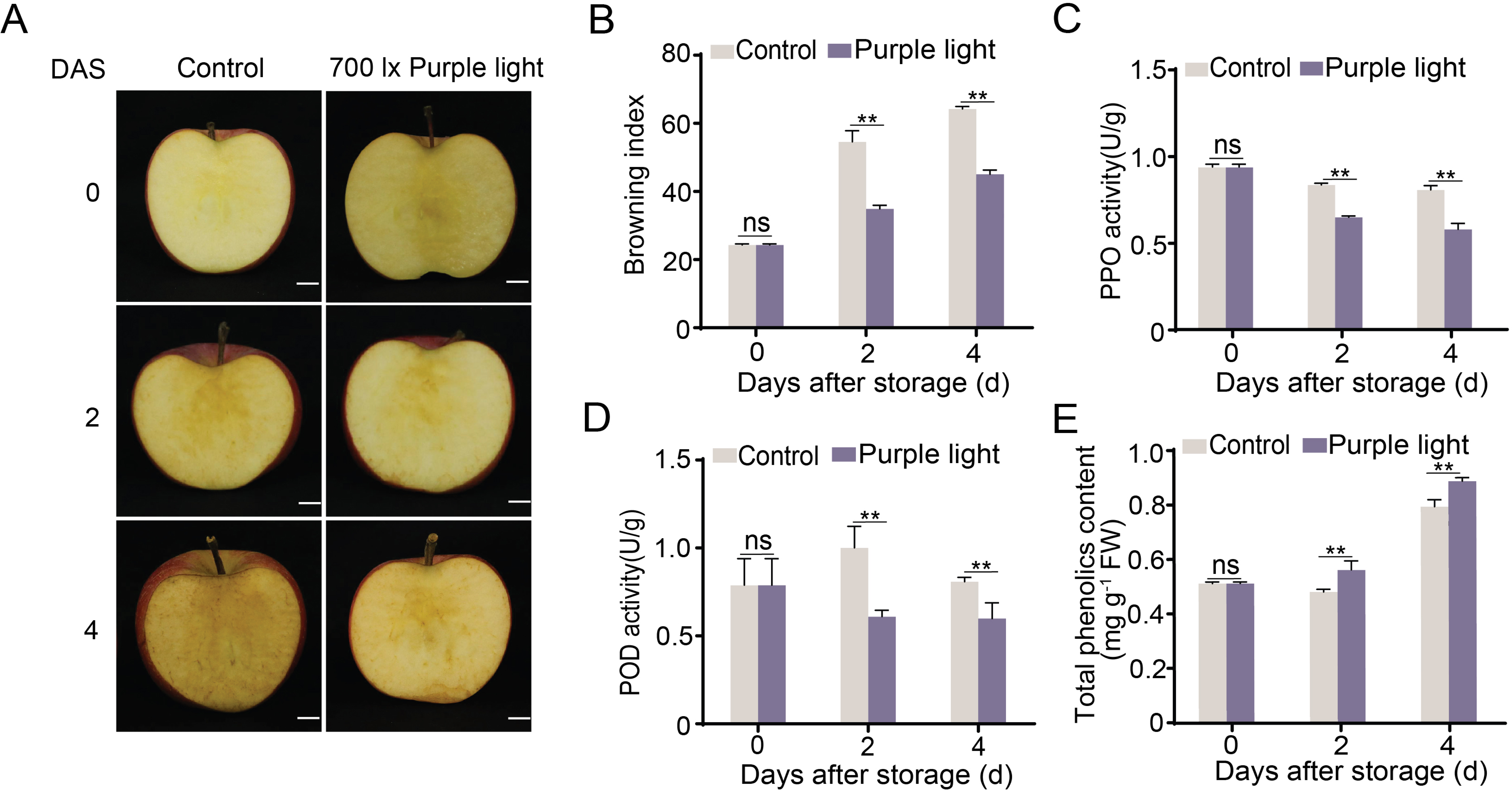
Purple light inhibits browning-related gene enzyme activity and increased total phenolic content in fresh-cut ‘Fuji’ apple. **(A)** ‘Fuji’ apples were harvested in 2020, 180 DAFB. Apple slices were treated with LED purple light at 700 lx in a 10 °C atmosphere for 4 d, and samples collected every 2 d. Samples were stored in the dark as a control. DAS: days after storage. Bars: 1 cm. **(B)** Browning index for control and purple light-treated samples. **(C)** PPO activity in control and purple light-treated samples. **(D)** POD activity in control and purple light-treated samples, replicated thrice. **(E)** Total phenolic content in control and purple light-treated samples. Data represent means ± SE. Asterisks indicate significant differences (*, P< 0.05; **, P < 0.01, Student’s *t-*test); ns, no significant difference.

To investigate the effect of purple light on browning, the optimal intensity of purple light was first determined by applying 700 lx, 1000 lx, and 1500 lx purple light to ‘Fuji’ apple slices, with results showing similar effects for all intensities; all decreased the browning index and inhibited browning (Figure S2A-F). To consider energy usage, 700 lx was therefore selected as the optimal light intensity. No difference was observed in the browning index of the controls, ‘Fuji’ apple that were subjected to natural light and darkness, while treatment with 700 lx purple light treatment led to a significant reduction in the browning index (Figure S2G, H). We therefore conclude that purple light at 700 lx can play an important role in inhibiting browning in fresh-cut apple.

PPO and POD directly affect the browning of freshly cut fruit. The PPO and POD activities in ‘Fuji’ apple was therefore examined, with results showing a decrease in the PPO activity during storage, both for apple treated with purple light and those tested under control conditions; however, lower PPO activity was observed over the entire storage period for apple treated with purple light as compared to the control (Figure 1C). Similar trends were observed in the POD and PPO activities, with direct inhibition of the POD activity observed during storage under purple light (Figure 1D). Phenolic compounds also play important roles in enzymatic browning, and treatment with purple light was observed to increase the phenolic content during storage (Figure 1E). Similar results were observed for ‘Hanfu’ and ‘Lvshuai’ (Figure S3; Figure S4).

Because the activities of PPO and POD were decreases and the content of phenolics were increases, and PAL is a key gene for phenolic synthesis, we speculated that purple light affects the gene expression of *MdPPO*, *MdPOD* and *MdPAL*. The associated genes *MdPPO*, *MdPOD*, and *MdPAL* were identified from apple genome, with *MdPPO* and *MdPOD* significantly decreased (Figure 2A, B), and *MdPAL* considerably enhanced (Figure 2C) by treatment with purple light. These results suggest that *MdPPO*, *MdPOD,* and *MdPAL* play important roles in browning.

**Figure 2.**
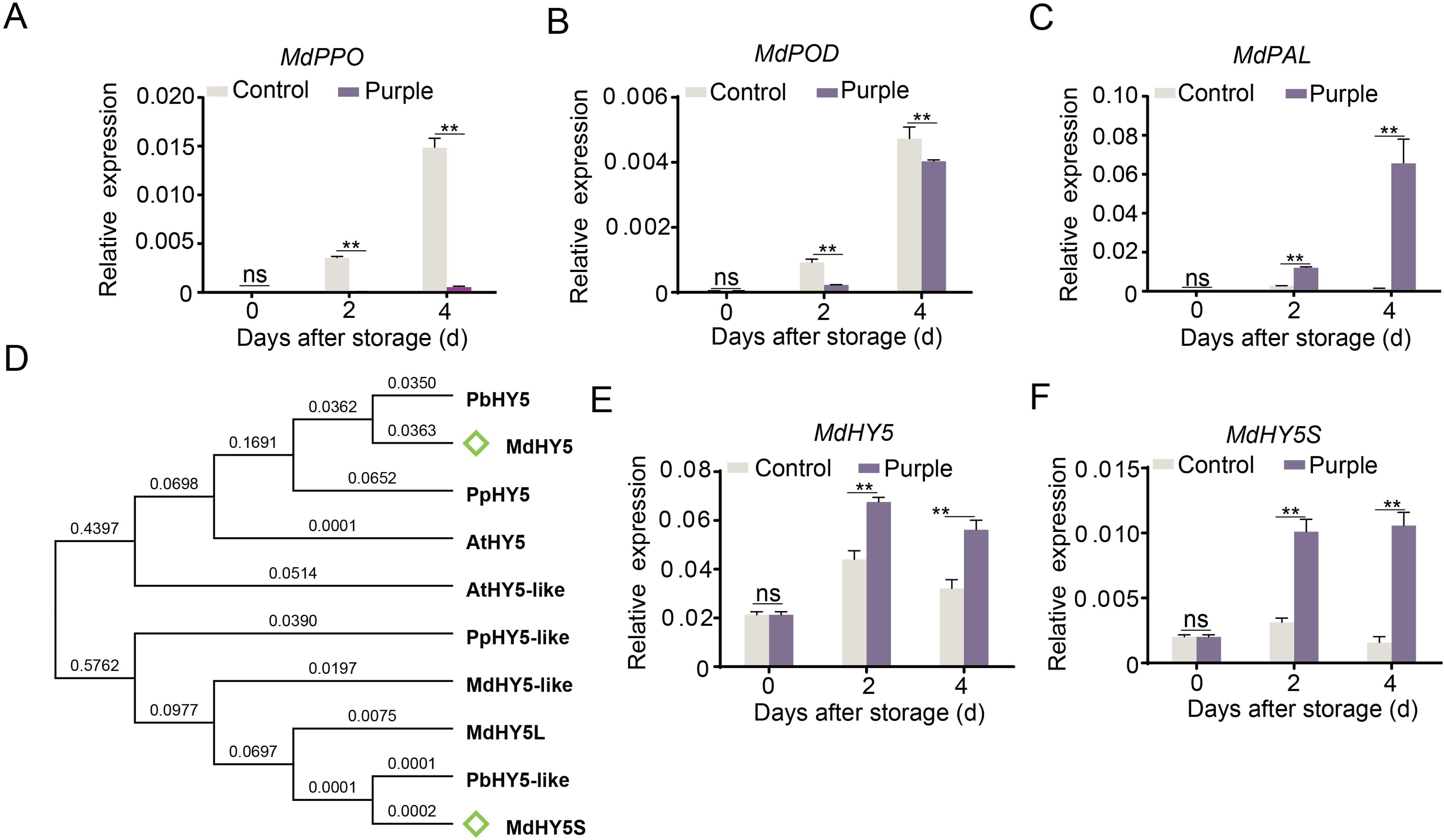
Gene expression influenced by purple light in fresh-cut apple. **(A)** *MdPPO e*xpression in purple light-treated samples was measured. **(B)** *MdPOD* expression level. **(C)** *MdPAL* expression level. **(D)** Phylogenetic tree analysis of HY5 during different species. **(E)** *MdHY5* expression level. **(F)** *MdHY5S* expression level, replicated thrice. Data represent means ± SE. Asterisks indicate significant differences (*, P< 0.05; **, P < 0.01, Student’s *t-*test); ns, no significant difference.

### *MdHY5* and *MdHY5S* are essential for purple light inhibition of browning in apple

HY5 is a key transcription factor in the light signaling pathway. To understand the role of HY5 in browning, four *MdHY5* genes were identified in the apple genome, with investigation of the phylogenetic tree indicating that they are closely related and have similar biological functions (Figure 2D). All four MdHY5 genes were found to be significantly upregulated by purple light (Figure S5). *MdHY5* and *MdHY5S,* with the highest expression, were selected as research objects (Fig 2E, F). To investigate the functions of *MdHY5* and *MdHY5S*, a transient expression assay was conducted using apple slices, with partial *MdHY5* and *MdHY5S* CDS (1–300 bp,1–300 bp) ligated into the pTRV2 vector to generate silencing vectors for *MdHY5* and *MdHY5S* (MdHY5-AN and MdHY5S-AN). The vectors were then transformed separately into *Agrobacterium tumefaciens* and the bacteria left to infect the apple slices over 4 d under purple light at 10 °C. An empty pTRV vector under purple light was used as a control. No MdHY5-AN and MdHY5S-AN fruit was placed in the dark during treatment because HY5 is degraded by COP1 under dark conditions. The results showed that purple light irradiation no longer inhibited the browning of fresh-cut apple after MdHY5 and MdHY5S were silenced, and the browning index was considerably higher than that of the control (Figure 3A, B, D, E). RT-qPCR results showed that the expression levels of *MdHY5* and *MdHY5S* were lower in MdHY5-AN and MdHY5S-AN fruit under purple light treatment than the control, while the expression levels of *MdPPO* and *MdPOD* were also higher and the expression level of *MdPAL* lower in the MdHY5-AN and MdHY5S-AN fruit as compared to the control (Figure 3C, F). These results demonstrate the important roles of *MdHY5* and *MdHY5S* in the browning process.

**Figure 3.**
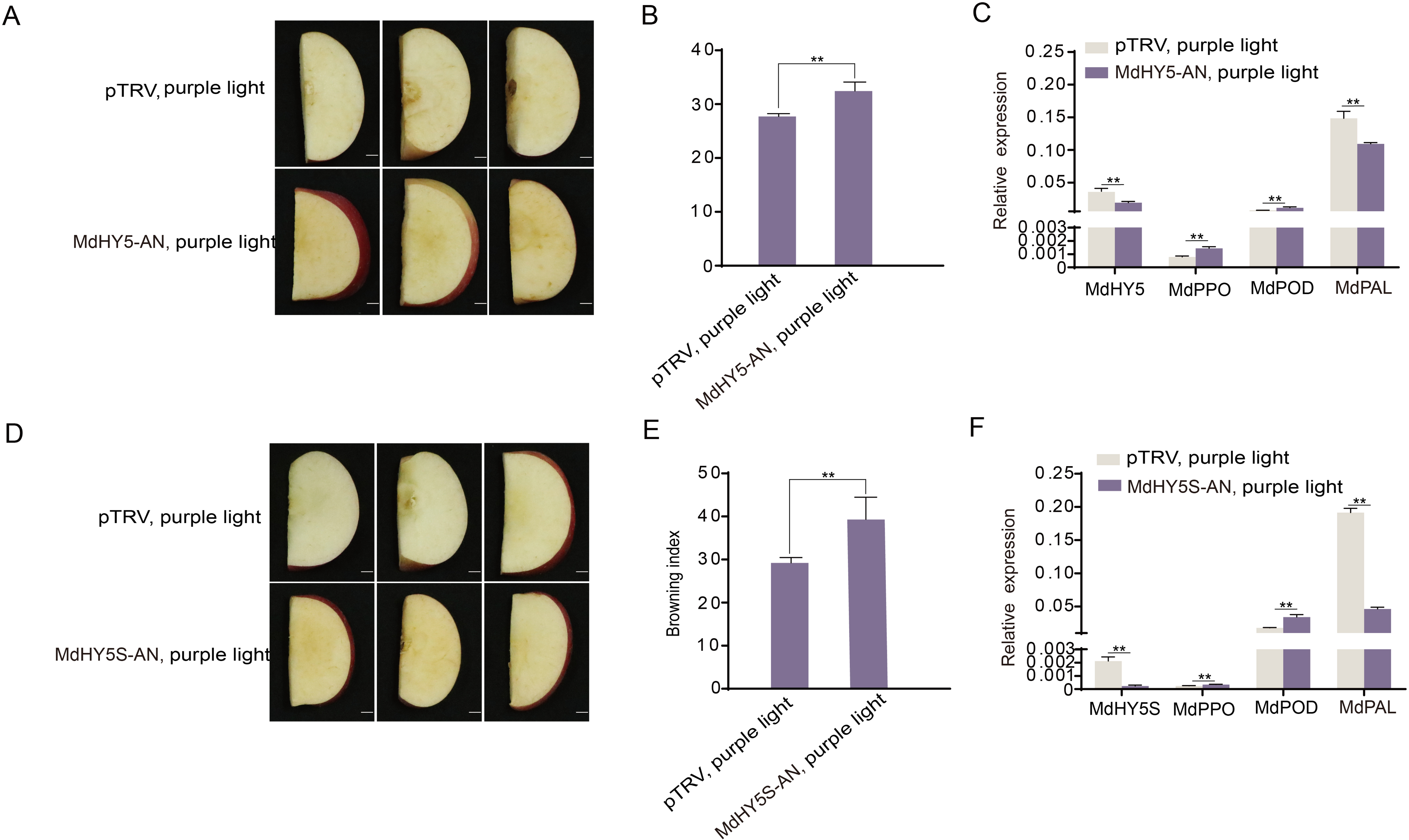
*MdHY5* and *MdHY5S* are required for purple light-induced fresh-cut fruit browning. **(A)** *MdHY5* was silenced in apple slices by transient infection, producing MdHY5-AN fruit that were then treated with purple light for 4 d. Empty vector-treated fruit (pTRV) were placed under purple light for 4 d as a control. Scale bars, 1 cm. **(B)** Browning index in MdHY5-AN fruit and control after purple light for 4 d. **(C)** *MdHY5, MdPPO, MdPOD, and MdPAL* expression in MdHY5-AN and control fruit, with qRT-PCR confirming successful infiltration. **(D)** Transient infection was used to silence *MdHY5S* in apple slices, after which the resulting MdHY5S-AN fruit was treated with purple light for 4 d. Empty vector-treated fruit (pTRV) was also placed under purple light for 4 d as a control. Scale bars, 1 cm. **(E)** Browning index in MdHY5S-AN fruit and control measured over 4 d by purple light. **(F)** Expression levels of *MdHY5S, MdPPO, MdPOD, and MdPAL* were evaluated in MdHY5S-AN and control fruit by qRT-PCR, confirming successful infiltration. The experiment was performed independently in three biological replicates. Data represent means ± SE. Asterisks indicate significant differences (*, P< 0.05; **, P < 0.01, Student’s *t-*test); ns, no significant difference.

To analyze their subcellular localization, *MdHY5* and *MdHY5S* CDS were ligated into pRI101 vector contain a green fluorescent protein (GFP) tag and overexpressed in tobacco leaves (*Nicotiana benthamiana*). Microscopy clearly showed that the MdHY5-GFP and MdHY5S-GFP proteins were localized to the nucleus (Figure S6).

### The binding of MdHY5 and MdHY5S to *MdPPO, MdPOD,* and *MdPAL* promoters and transcription regulation

Silencing *MdHY5* and *MdHY5S* in apple was found to induce *MdPPO* and *MdPOD* expression and decrease *MdPAL* expression. We therefore hypothesized that MdHY5 and MdHY5S regulate the transcription of *MdPPO*, *MdPOD,* and *MdPAL*. Promoter analysis indicated the presence of four G-boxes (light-response elements) in the *MdPPO* promoter, three in the *MdPOD* promoter, and six in the *MdPAL* promoter. We first determined whether MdHY5 and MdHY5S could bind to the *MdPPO*, *MdPOD,* and *MdPAL* promoters using a yeast one-hybrid (Y1H) assay, with results showing that full-length MdHY5 and MdHY5S could bind to the *MdPPO* and *MdPAL* promoters (Figure 4A; Figure 6A), while the *MdPOD* promoter was not inhibited by ABA^1000^ (Figure S7). The first fragment on the promoter that contained G-box binding sites was selected as a probe for the Electrophoretic mobility shift assay (EMSA) experiments. The results showed that MdHY5 and MdHY5S bound to the *MdPPO*, *MdPOD,* and *MdPAL* promoters via the first binding sites, and that the addition of unlabeled competing probes led to weakening of the binding bands. Treatment with thermal mutation probes was found to inhibit the binding of MdHY5 and MdHY5S with the promoters (Figure 4B; Figure 5A; Figure 6B). These results demonstrate that MdHY5 and MdHY5S can bind to the *MdPPO*, *MdPOD,* and *MdPAL* promoters in vitro. To verify whether MdHY5 and MdHY5S could bind to the *MdPPO*, *MdPOD,* and *MdPAL* promoters in vivo, *MdHY5* and *MdHY5S* were ligated into the upstream of a FLAG tag in pRI101 vector to overexpress *MdHY5* and *MdHY5S* in apple calli. Chromatin immunoprecipitation (ChIP)-qPCR assay indicated enrichment of the *MdPPO*, *MdPOD,* and *MdPAL* promoters in *MdHY5* and *MdHY5S* overexpressed apple calli (Figure 4C; Figure 5B; Figure 6C).

**Figure 4.**
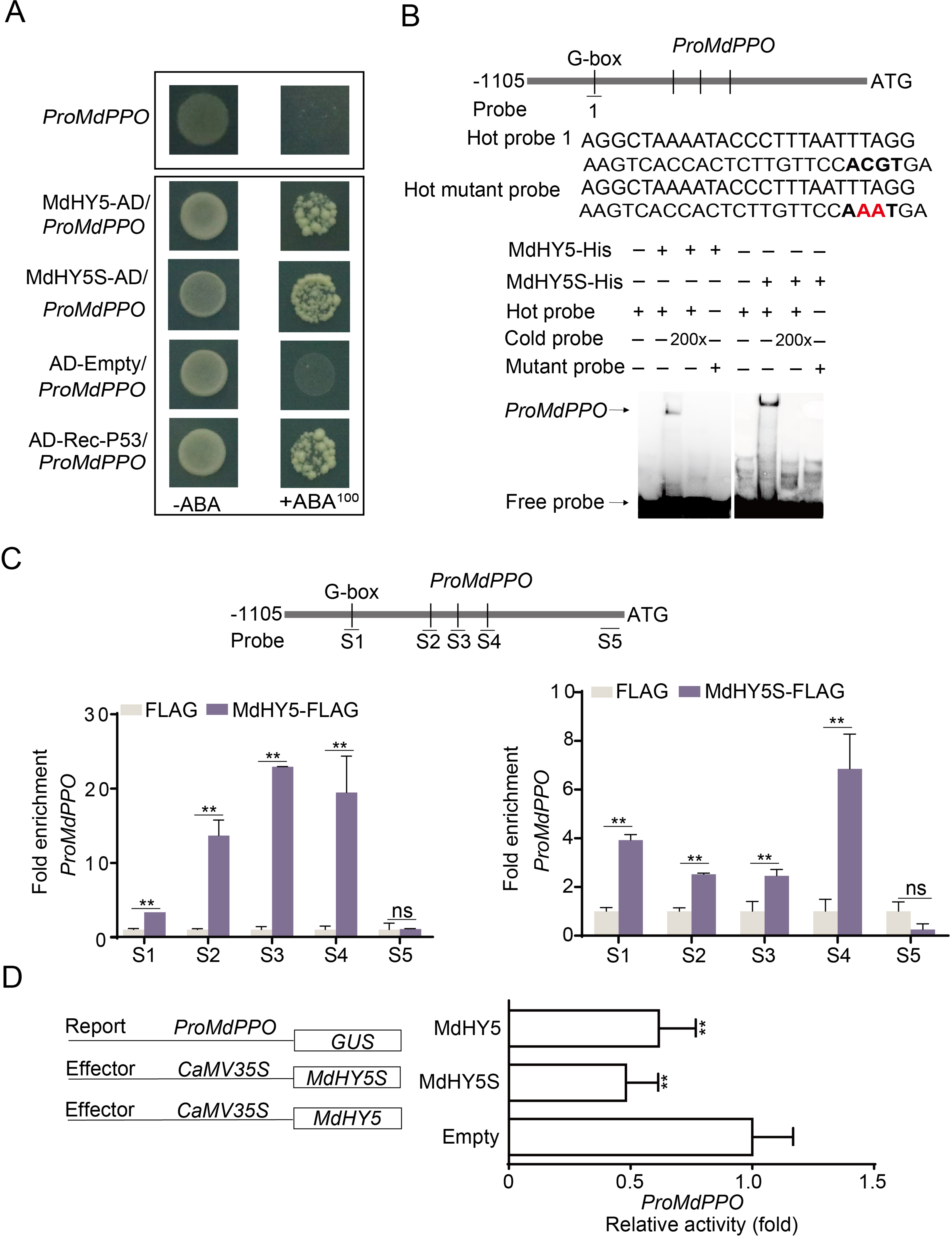
*MdHY5* and *MdHY5S* negatively regulated *MdPPO*. **(A)** Y1H analysis showed the MdHY5 and MdHY5S bound to the *MdPPO* promoter. P53 was acted positive control. Empty vector pGADT7 (AD) was acted negative control. **(B)** EMSA analysis showed that MdHY5 and MdHY5S bound to the *MdPPO* promoter fragment with G-box. Hot probe was designed to contain biotin labeling, and unlabeled promoter fragment as the cold competitive probe (at a concentration 200 x). Hot probe with two nucleotide mutation were used as the mutation probe. Mutation sites were marked in red. MdHY5-His protein and MdHY5S-His were purified. The probe sequence was shown above. The ‘‘+’’ or ‘‘-’’ meant presence and absence. **(C)** ChIP-PCR analysis showed that MdHY5 and MdHY5S bound to the *MdPPO* promoter in vivo. The FLAG antibody was used to verify the binding of MdHY5 and MdHY5S to the *MdPPO* promote and result was detected by quantitative(qPCR). Fruit calli with FLAG tag alone was used as a negative control. The experiment was performed independently in three biological replicates. **(D)** GUS activation assay illuminated that MdHY5 and MdHY5S negatively regulated the *MdPPO* promoter. The experiment was performed independently in three biological replicates. Data represent means ± SE. Asterisks indicate significant differences (*, P< 0.05; **, P < 0.01, Student’s *t-*test); ns, no significant difference.

**Figure 5.**
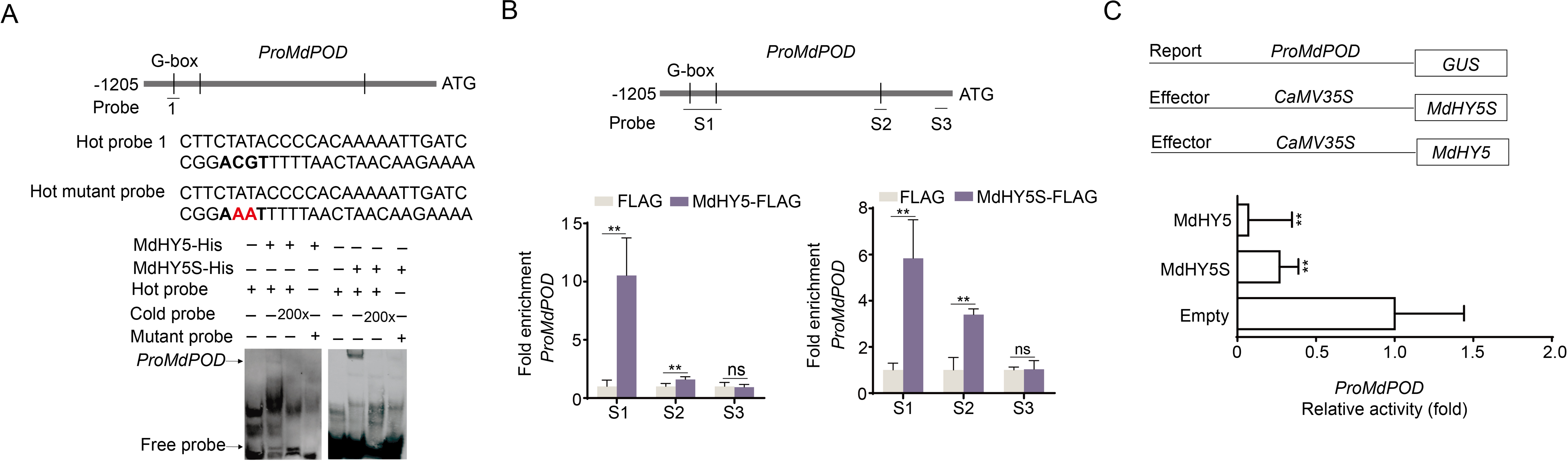
*MdHY5* and *MdHY5S* negatively regulated *MdPOD*. **(A)** EMSA analysis showed MdHY5 and MdHY5S bound to *MdPOD* promoter fragment with G-box. Hot probe was designed to contain biotin labeling, and unlabeled promoter fragment as the cold competitive probe (at a concentration 200 x). Hot probe with two nucleotide mutation were used as the mutation probe. Mutation sites were marked in red. MdHY5-His protein and MdHY5S-His were purified. The probe sequence was shown above. The ‘‘+’’ or ‘‘-’’ meant presence and absence. **(B)** ChIP-PCR analysis showed that MdHY5 and MdHY5S bound to the *MdPOD* promoter in vivo. The FLAG antibody was used to verify the binding of MdHY5 and MdHY5S to the *MdPOD* promote and result was detected by quantitative(qPCR). Fruit calli with FLAG tag alone was used as a negative control. The experiment was performed independently in three biological replicates. **(C)** GUS activation assay illuminated MdHY5 and MdHY5S negatively regulate the *MdPOD* promoter. The experiment was performed independently in three biological replicates. Data represent means ± SE. Asterisks indicate significant differences (*, P< 0.05; **, P < 0.01, Student’s *t-*test); ns, no significant difference.

**Figure 6.**
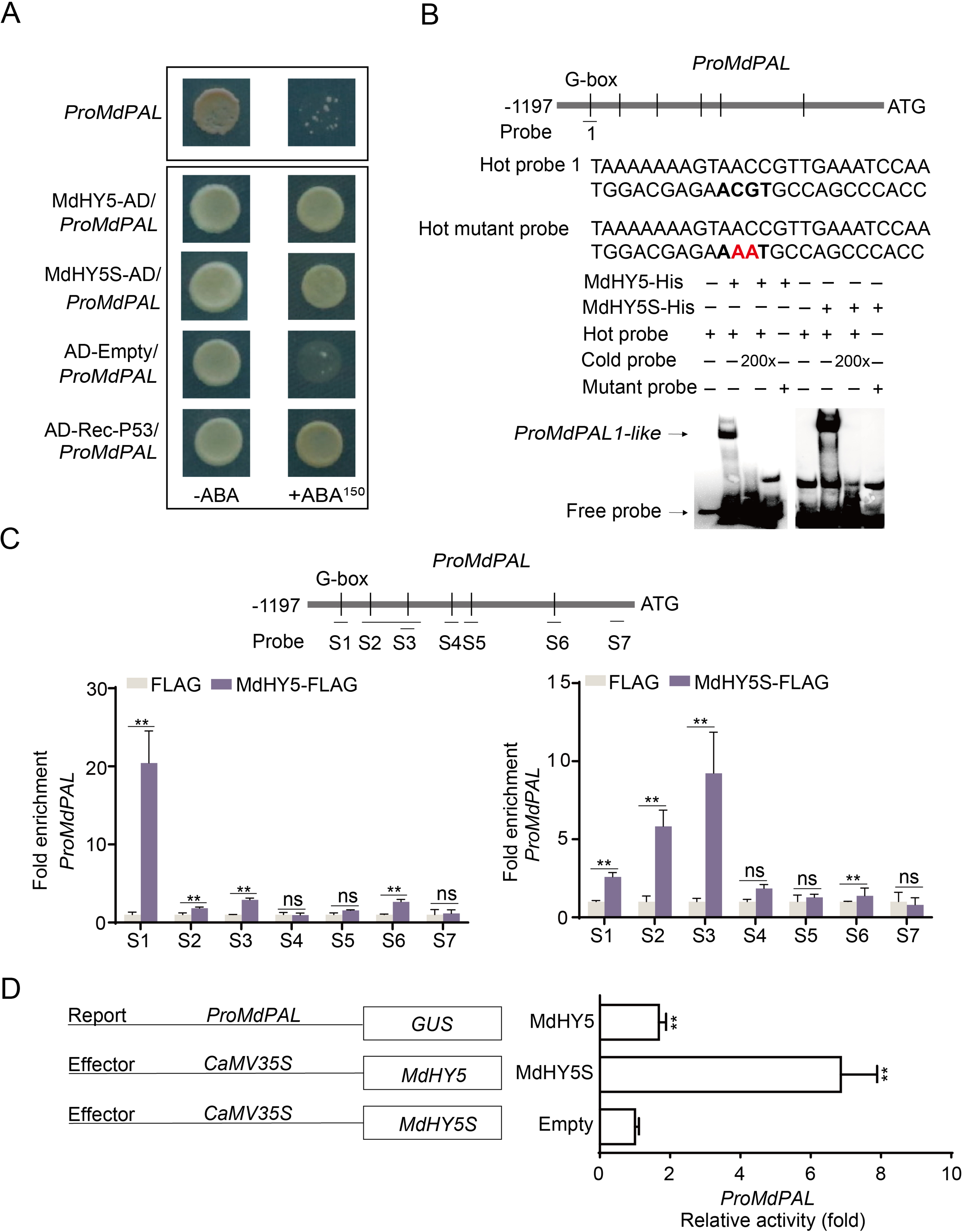
*MdHY5* and *MdHY5S* positively regulated *MdPAL*. **(A)** Y1H analysis showed MdHY5 and MdHY5S bound to the *MdPAL* promoter. P53 was acted positive control. Empty vector pGADT7 (AD) was acted negative control. **(B)** EMSA analysis showed that MdHY5 and MdHY5S bound to the *MdPAL* promoter fragment with G-box. Hot probe was designed to contain biotin labeling, and unlabeled promoter fragment as the cold competitive probe (at a concentration 200 x). Hot probe with two nucleotide mutation were used as the mutation probe. Mutation sites were marked in red. MdHY5-His protein and MdHY5S-His were purified. The probe sequence was shown above. The ‘‘+’’ or ‘‘-’’ meant presence and absence. **(C)** ChIP-PCR analysis showed MdHY5 and MdHY5S bound to the *MdPAL* promoter in vivo. The FLAG antibody was used to verify the binding of MdHY5 and MdHY5S to the *MdPAL* promote and result was detected by quantitative(qPCR). Fruit calli with FLAG tag alone was used as a negative control. The experiment was performed independently in three biological replicates. **(D)** GUS activation assay illuminated MdHY5 and MdHY5S negatively regulated the *MdPAL* promoter. The experiment was performed independently in three biological replicates. Data represent means ± SE. Asterisks indicate significant differences (*, P< 0.05; **, P < 0.01, Student’s *t-*test); ns, no significant difference.

Next, a β-glucuronidase (GUS) activation assay was used to verify the regulation of *MdPPO*, *MdPOD,* and *MdPAL* by MdHY5 and MdHY5S in *Nicotiana benthamiana* leaves. We found that the GUS activity that co-expressed *pro35S:MdHY5* with *proMdPPO:GUS* and *proMdPOD:GUS* was significantly lower than that of the control, which is similar to the co-expression of *pro35S:MdHY5S* with *proMdPPO:GUS* and *proMdPOD:GUS,* and suggests that MdHY5 and *MdHY5S* are transcriptional inactivators for *MdPPO* and *MdPOD* (Figure 4D; Figure 5C). Collectively, these results suggest that MdHY5 and MdHY5S can bind to the *MdPPO* and *MdPOD* promoters both in vitro and in vivo, inhibiting transcription. The GUS activities of *pro35S:MdHY5* and *pro35S:MdHY5S* with *proMdPAL:GUS* were significantly higher than those in the control, suggesting that MdHY5 and MdHY5S induce *MdPAL* expression as transcriptional activators (Figure 6D). These results suggest that MdHY5 and MdHY5S bind to the *MdPAL* promoter, promoting its transcription both in vitro and in vivo.

### The promotion of MdHY5 and MdHY5S expression via interaction with corresponding promoters

To explore the relationship between *MdHY5* and *MdHY5S*, two-hybrid yeast experiments were performed. The result showed no interaction between MdHY5 and MdHY5S (Figure S8). To clarify the upstream and downstream relationships of *MdHY5* and *MdHY5S*, the *MdHY5* and *MdHY5S* promoters were analyzed, with results showing that the *MdHY5* promoter contains three and the *MdHY5S* promoter four G-boxes. Y1H and EMSA assays were therefore conducted, with results suggesting that MdHY5 binds to the *MdHY5S* promoter and MdHY5S binds to the *MdHY5* promoter in vitro (Figure 7A-D). ChIP-qPCR results showed that MdHY5 leads to *MdHY5S* promoter fragment enrichment as compared to the control. Similarly, MdHY5S was enriched in *MdHY5* promoter fragments (Figure 7E, F). These results show that MdHY5 and MdHY5S bind to their corresponding promoters both in vitro and in vivo.

**Figure 7.**
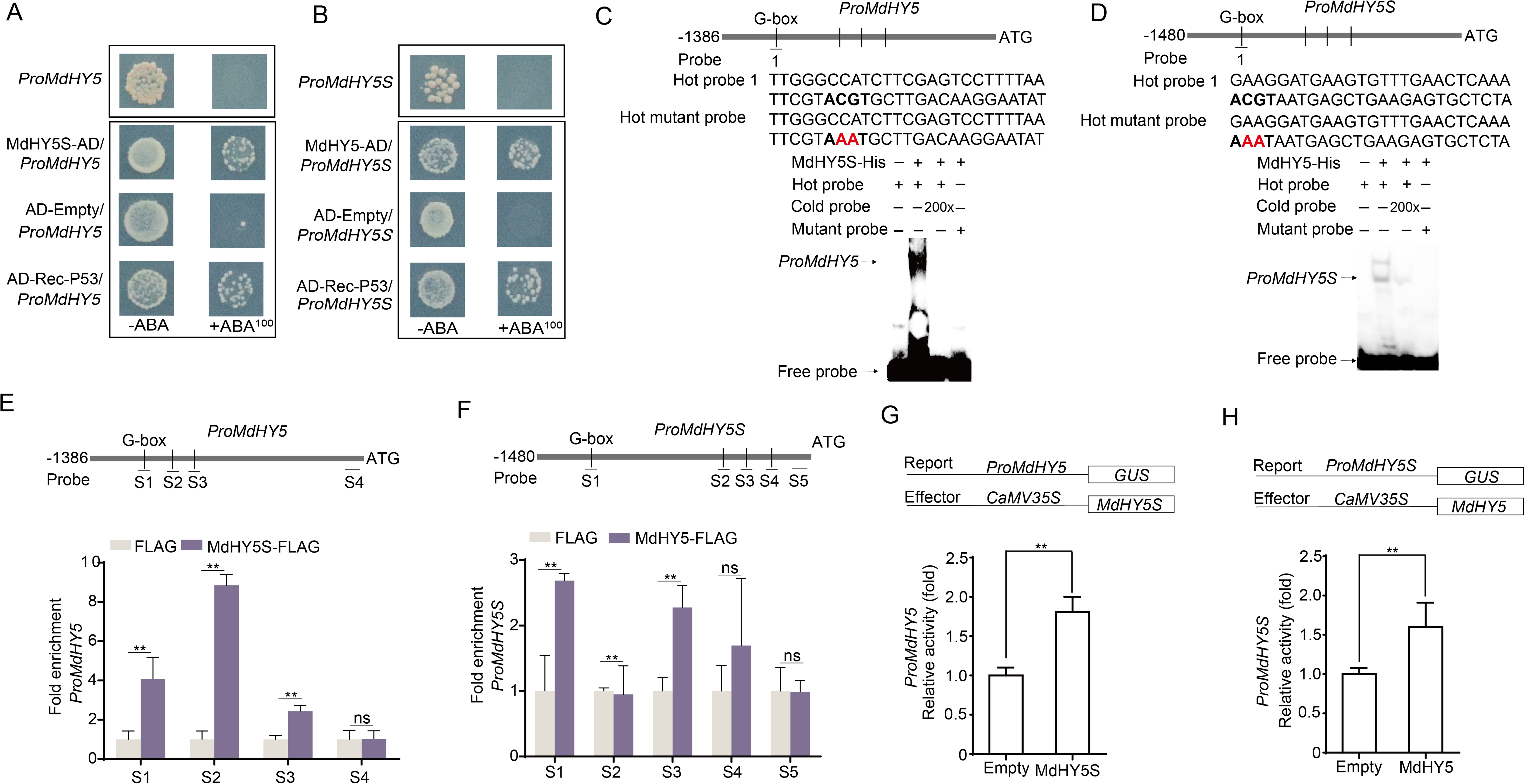
MdHY5 and MdHY5S actively regulated each other’s promoter. **(A, B)** Y1H analysis showed MdHY5 and MdHY5S bound to each other’s promoter. P53 was acted positive control. Empty vector pGADT7 (AD) was acted negative control. **(C, D)** EMSA analysis showed MdHY5 and MdHY5S bound to each other’s promoter fragment with G-box. Hot probe was designed to contain biotin labeling, and unlabeled promoter fragment as the cold competitive probe (at a concentration 200 x). Hot probe with two nucleotide mutation were used as the mutation probe. Mutation sites were marked in red. MdHY5-His protein and MdHY5S-His were purified. The probe sequence was shown above. The ‘‘+’’ or ‘‘-’’ meant presence and absence. **(E, F)** ChIP-PCR analysis showed MdHY5 and MdHY5S bound to each other’s promoter in vivo. The FLAG antibody was used to verify the binding of MdHY5 and MdHY5S to each other promoter and result was detected by quantitative(qPCR). Fruit calli with FLAG tag alone was used as a negative control. The experiment was performed independently in three biological replicates. **(G, H**) GUS activation assay showed MdHY5 positively regulates the *MdHY5S* promoter and MdHY5S positively regulates the *MdHY5* promoter. The experiment was performed independently in three biological replicates. Data represent means ± SE. Asterisks indicate significant differences (*, P< 0.05; **, P < 0.01, Student’s *t-*test); ns, no significant difference.

We then co-expressed *pro35S:MdHY5* with *proMdHY5S* in *Nicotiana benthamiana* leaves and found higher GUS activity as compared to the control. Additionally, the GUS activity of the co-expressing *pro35S:MdHY5S* with *proMdHY5* in *Nicotiana benthamiana* leaves was also higher (Figure 7G, H). These results show that MdHY5 and MdHY5S positively regulate their corresponding promoter activities.

## Discussion

Light is involved in both the growth and development of plants (Huang et al. 2022; Liu et al. 2022). Several studies have described the effect of light on the quality of fruit, including tomatoes (Ntagkas et al. 2019), grapes (Azuma et al. 2019), strawberries (Xu et al. 2018), sweet cherries (Wang et al. 2023), peaches (Zhao et al. 2017), citrus (Lafuente et al. 2021), melons (Liu et al. 2022), and pears (Tao et al. 2018). However, only changes in the coloration and nutrient content have thus far been explored, and the effect that light has on browning in fresh-cut fruit remains unclear.

### The role of purple light in inhibiting browning in fresh-cut fruit

Light treatment is a simple, flexible, and cost-effective method, especially visible light, which is more convenient and pollution-free as compared to UV-C, UV-A, and UV-B (Lante et al. 2016; Li et al. 2022; Liu et al. 2023). We noticed that LED light sources of all different qualities reduced the browning index of fresh-cut apple and inhibited browning in ‘Fuji’ apple, with the effect of purple light the most obvious (Figure 1A, B; Figure S1A, B). Moreover, the browning rate did not change significantly under different light intensities (Figure S2), indicating that light quality rather than intensity is the important factor that affects fruit browning. Treatment with light of different qualities varieties showed that purple light had the best effect in inhibiting browning in both ‘Hanfu’ and ‘Lvshuai’ (Figure S1C-F), suggesting that purple light influences browning in several fresh-cut apple varieties. However, this finding has not been mentioned in previous studies.

### The phenolics act as antioxidant in inhibiting browning in fresh-cut apple

The enzymatic browning of fruit and vegetables is a complex physiological and biochemical process that involves the oxidation of phenolic substances to quinones. Phenolics serve not only as oxidation substrates for enzymes, but also as bioactive compounds with antioxidant activity that have beneficial effects on human health. The biosynthesis of phenolic compounds in fruit and vegetables is induced by wounding stress, which inhibits oxidation (Li et al. 2017), with total phenolic content depending on the balance between biosynthesis and conversion. In other words, the observed phenolic content increase indicates that the synthesis rate is higher than the conversion rate and that phenolic compounds are being synthesized as a defense mechanism, while a decrease in the phenolic content would indicate that phenolic compounds have been converted into quinones (Fernando Reyes et al. 2007). In this study, the phenolic content increased following purple light irradiation, (Figure 1E), indicating that the light induced phenolic generation, with higher phenolic levels and a lower browning index observed as compared to the control (Figure 1B). This suggests that phenolic synthesis, rather than conversion, acts as a defense mechanism. The increase in phenolics not only inhibits the occurrence of browning, but also improves the quality of the fruit. A similar result was obtained for freshly cut pitaya. Hot air pretreatment suppresses the browning of fresh-cut pitaya fruit by regulating the phenylpropanoid pathway and ascorbate-glutathione cycle (Li et al. 2022). Cold plasma has also been observed to increase the phenolic content of fresh-cut mango (Yi et al. 2022).

### HY5 plays both positive and negative regulatory roles in inhibiting fresh-cut fruit browning

HY5 is an important transcription factor in the light signaling pathway that affects both the growth and development of plants, fruit coloration, and metabolite synthesis (Job et al. 2022; Sharma et al. 2023); however, the role of HY5 in these processes remains unclear. In this study, silencing MdHY5 and MdHY5S in fruit stored under purple light resulted in an increase in the expression levels of the browning-related genes *MdPPO* and *MdPOD*, and a decrease in the expression level of the phenolic synthesis gene *MdPAL* (Figure 3C, F). The results confirm that MdHY5 and MdHY5S play important roles in the inhibition of browning by purple light. Moreover, we found that MdHY5 and MdHY5S can bind to *MdPPO* and *MdPOD* promoters, negatively regulating their transcription and inhibiting browning, while at the same time, MdHY5 and MdHY5S increase the antioxidant phenolic content by positively regulating *MdPAL* transcription, again inhibiting browning (Figure 4; Figure 5; Figure 6). In addition, MdHY5 and MdHY5S were found to interact with their corresponding promoters to form a positive feedback transcriptional regulatory loop (Figure 7). Previous studies have shown that HY5 can inhibit the synthesis of ethylene as a negative regulator (Wang et al. 2023) and promote coloration as a positive regulator (Wang et al. 2021). In this study, we found that MdHY5 and MdHY5S which are induced by purple light, acts as both a positive and negative regulator. This was not mentioned in previous studies of HY5. Similarly, MYB306-like factor binds to the MYB17 promoter to initiate transcription and to the MdDFR promoter, inhibiting transcription in apple (Wang et al. 2022).

Finally, a model was developed suggesting how purple light regulates browning and phenolic sythesis in freshly cut apple. The application of purple light irradiation led to inhibition of the COP1 activity and translocation from the nucleus to the cytoplasm. *MdHY5* and *MdHY5S* expression was activated by purple light. *MdHY5* and *MdHY5S* inhibits transcription of the browning genes *MdPPO* and *MdPOD* by directly binding to the promoter, whereas MdHY5 and MdHY5S bind to the *MdPAL* promoter, increasing both the phenolic content and the antioxidant activity. Moreover, *MdHY5* and *MdHY5S* bind to each other’s promoters, creating a positive feedback loop and inhibiting browning. Light cessation led to the degradation of MdHY5 and MdHY5S by COP1 and decreased *MdHY5* and *MdHY5S* expression, weakening the transcription of *MdPPO*, *MdPOD,* and *MdPAL* and resulting in browning (Figure 8).

**Figure 8.**
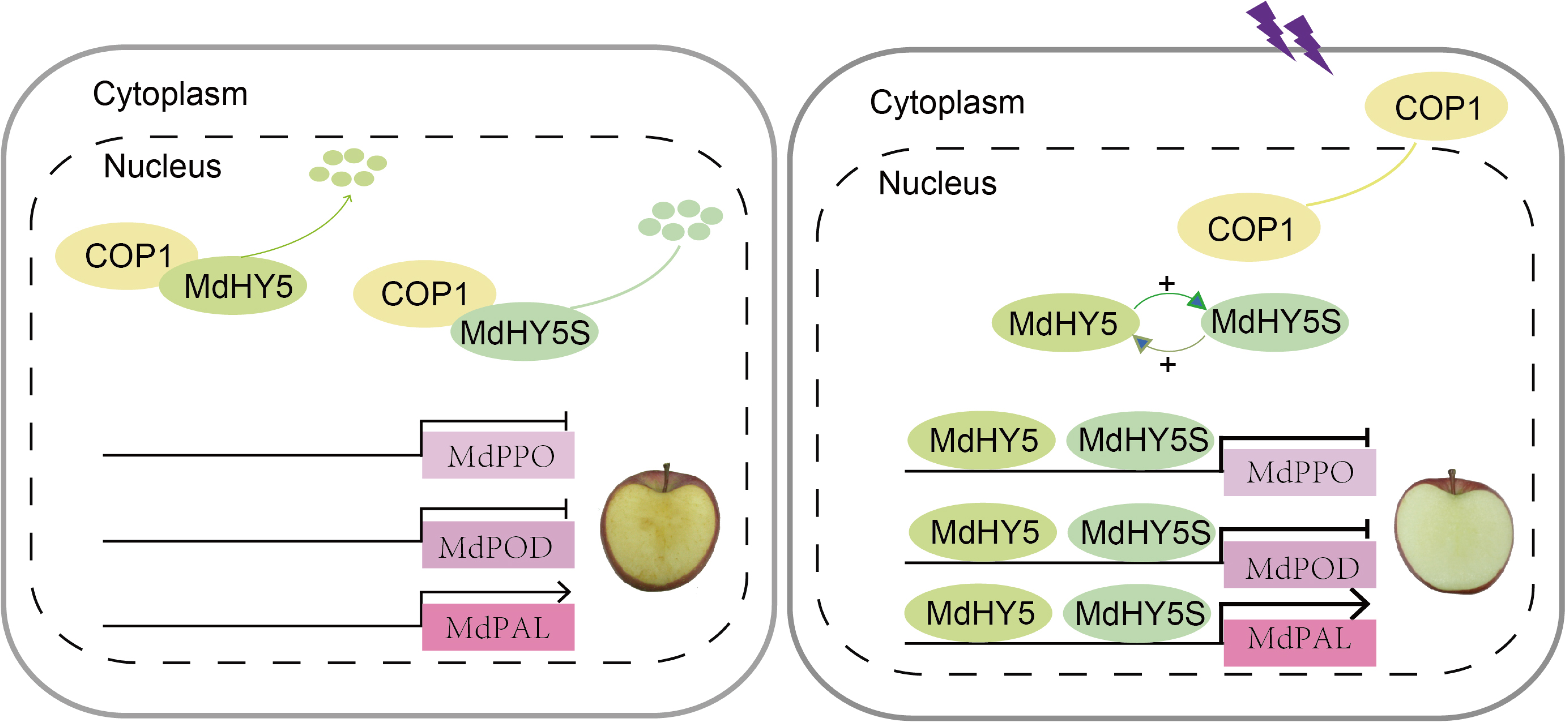
Model showing the molecular mechanism by which purple light inhibits browning in fresh-cut apple and promotes phenolic sythensis. After exposure to purple light, COP1 activity was inhibited and translocated from the nucleus to the cytoplasm; MdHY5 and MdHY5S promoted each other’s expression by combining with each other’s promoters. Then, *MdHY5* and *MdHY5S* inhibited the browning gene *MdPPO*, *MdPOD* transcription occurred with a direct binding promoter, and MdHY5 and MdHY5S bound to the *MdPAL* promoter to increase the phenolic content and antioxidant activity, thereby inhibiting browning.

## Materials and Methods

### Plant material and treatments

‘Fuji’ apple (*Malus x domestica* Borkh.) were obtained from the experimental farm at the Liaoning Pomology Institute (Xiongyue, China) in 2020. Fruit were collected with the commercial harvest (180 DAFB) and transferred immediately to the laboratory, where they were stored in a refrigerator for testing. Refrigerated apple were cut into slices and divided into nine groups (25 slices per group), of which the first eight groups were sequentially treated with red, orange, yellow, green, indigo, blue, purple, and white LED light, and the ninth was placed directly in the dark as a control. The light intensity was set to 700 × and all fruit were placed at 10 °C for 4 d. Ten slices were then removed every 2 d, frozen in liquid nitrogen, and stored at −80 °C for further use.

### Measurement of Browning Index

The flesh color of fresh-cut fruit were measured using a chroma meter (chroma meter CR-400) and L*, a*, and b* values for the fruit surface recorded. The browning index was calculated using equations 1 and 2 (Zheng et al. 2019):

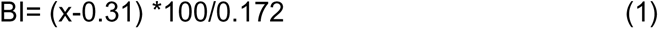

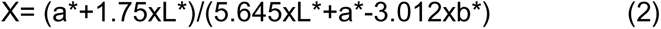

### Measurement of PPO and POD Activities

The PPO and POD activities were extracted according to Simões et al. (2015).

For PPO activity, 500 μL of crude extract was added to 500 μL of 0.2 mol/L catechins and 500 μL of phosphoric acid buffer (pH 6.5), and a UV/VIS spectrophotometer used to measure the absorbance at 420 nm.

For POD activity, 500 µL sodium phosphate buffer (0.2 mol L^-1^, pH 6.0) was added to 100 µL enzyme extract, 200 µL guaiacol (0.5 %), and 200 µL H_2_O_2_ at 0.08 % (v/v) and measured using UV/VIS spectrophotometry.

One unit of PPO and POD activity was defined as a change of 0.01 per min. And they are presented as U/g fresh weight.

### Measurement of total phenolic content

Apple powder (0.5 g) was extracted by homogenization with 5 mL of 80 % (v/v) methanol and centrifuged at 10 000 g for 20 min. The total phenolic content was described by Fan et al. (2018).

### RNA extraction and gene expression analysis

Total RNA was extracted reference Li et al. (2016). Gene expression was performed by reverse transcription quantitative PCR (RT-qPCR) (Li et al. 2017). Apple ACTIN (EB136338) was used as an internal reference gene.

### Phylogenetic tree construction

The MEGA 11(Tamura et al. 2021) program was performed to construct phylogenetic. NCBI accession numbers are as follows: *AtHY5* (*Arabidopsis thaliana*; BT025519.1), *AtHY5-like* (*Arabidopsis thaliana*, NP_001330553.1), PpHY5 (*Prunus persica*; XP_020411091.1), *PbHY5* (*Pyrus x bretschneideri*; QGP73826.1), *PbHY5-like* (*Pyrus x bretschneideri*; XM_009355182.3), *PpHY5-like* (*Prunus persica*; XP_007223521.1), *MdHY5* (*Malus × domestica*; AB710143), *MdHY5* (*Malus × domestica*; XM_008371356), *MdHY5* (*Malus × domestica*; MD04G005220), *MdHY5* (*Malus × domestica*; NM_001293960).

### Subcellular localization of MdHY5 and MdHY5S

*MdHY5* and *MdHY5S* were ligated separately into pRI101 vector contain GFP tag (*35S:GFP-MdHY5, 35S:GFP-MdHY5S*). MdHY5-GFP and MdHY5S-GFP injection buffers were injected into tobacco leaves and GFP used as a control. A confocal microscope was used to observe the fluorescence after three days, and NF-YA4-mCherry (Zhang et al. 2019) was used as a nuclear marker.

### *Agrobacterium*-mediated infiltration

To overexpress *MdHY5* and *MdHY5S* in apple calli (Orin), full-length *MdHY5* and *MdHY5S* were ligated upstream of 3xFLAG in the pRI101 vector with *35S* to form *Pro35S:MdHY5-FLAG* and *Pro35S:MdHY5S-FLAG*. To silence *MdHY5* and *MdHY5S* in apple, partial *MdHY5* and *MdHY5S* CDS (3– 303 bp,1–300 bp) were ligated onto the pTRV2 vector to form MdHY5-AN and MdHY5S-AN vectors. Apple slices infection refer to the method of Wei et al. (2021), with slight modification. In short, fresh-cut apple slices were placed in the injection buffer and incubated for 1 h, and the injection solution infiltrated into apple samples by vacuuming, after which samples were immediately placed in the light incubator at 10 °C and treated with purple light for 4 d. To infect apple calli, calli were resuspended in 50 ml MS liquid medium with 1 mM acetosyringone for 20 min, collected, spread on MS solid medium, and cultivated for 3 d.

### Y1H assay

*MdHY5* and *MdHY5S* were cloned into a pGADT7 vector. *MdPPO*, *MdPOD*, and *MdPAL* promoter fragments were ligated onto pAbAi vector. The Y1H assay reference Li et al. (2016).

### Electrophoretic mobility shift assay (EMSA)

*MdHY5* and *MdHY5S* were ligated into His-tagged vector and transformed into competent Escherichia coli BL21 (DE3) cells (TransGen Biotech). Proteins were then purified using the method described in Li (2017). For EMSA, a 3’-biotin-end-labeled double-stranded DNA probe was prepared. EMSA was performed using the Light Shift Chemiluminescence EMSA Kit, as described previously (Li et al. 2016).

### Chromatin immunoprecipitation (ChIP)-PCR analysis

*MdHY5* and *MdHY5S* were cloned into the pRI101 vector with a 3xFLAG tag. The CHIP assay was performed using a CHIP kit in accordance with the manufacturer’s instructions. The FLAG antibody was used to verify the binding of MdHY5 and MdHY5S to the *MdPPO*, *MdPOD*, and *MdPAL* promoters, and enrichment of the immunoprecipitates analyzed by PCR. The calli was infected three times and three ChIP assays were performed as three biological replicates.

### GUS activation assay

*MdPPO*, *MdPOD*, and *MdPAL* promoters were connected to the upstream region of the GUS reporter gene in the PBI121 vector as a reporter, and *MdHY5* and *MdHY5S* cloned into the pRI101 overexpression vector as effectors. Finally, the reporter and effector factors were co-injected into tobacco leaves. GUS activity was measured using the method in Li et al. (2016).

### Accession numbers

Sequence data from this study can be found in the Genome Database for Rosaceae (https://www.rosaceae.org) or the GenBank libraries under the accession numbers *MdActin* (EB136338), *MdPPO* (MD05G1319800), *MdPOD* (MD14G1010300), *MdPAL* (MD04G1096200), *MdHY5* (AB710143), and *MdHY5S* (XM_008371356).

## Author contributions

Aide Wang designed the experiments; Juntong Jin, Liyong Qi, Shuran Yang, Shurong Shen participated in experiments and analyzed data; Juntong Jin wrote the manuscript with inputs and guidance from Aide Wang and Hui yuan. All authors have read and approved the final manuscript.

## Acknowledgements

We thank editage (https://www.editage.cn/) for editing the manuscript.

